# Multiscale Adaptive Gabor Expansion (MAGE): Improved Detection of Transient Oscillatory Burst Amplitude and Phase

**DOI:** 10.1101/369116

**Authors:** Ryan T. Canolty, Thilo Womelsdorf

## Abstract

— Since Denis Gabor’s pioneering paper on the discrete Gabor Expansion (Gabor, 1946), time-frequency signal analysis has proven to be an important tool for many fields. In neurophysiology, time-frequency analysis has often been used to characterize and describe transient bursts in local field potential data. However, these transient bursts have a wide range of variable durations, suggesting that a time-frequency-scale dictionary composed of elementary signal “atoms” may prove useful to more accurately match recorded bursts. While overcomplete multiscale dictionaries are useful, generating a sparse code over such dictionaries is a difficult computational problem. Existing adaptive algorithms for discovering a sparse description are slow and computationally intensive. Here we describe the Multiscale Adaptive Gabor Expansion (MAGE), which uses an implicit dictionary of parametric time-frequency-scale Gabor atoms to perform fast parameter reassignment to accelerate discovery of a sparse decomposition. Using analytic expressions together with numerical computation, MAGE is a greedy pursuit algorithm similar to Matching Pursuit, restricted to a dictionary of multiscale Gaussian-envelope Gabor atoms. MAGE parameter reassignment is robust in the presence of moderate noise. By expressing a unknown signal as a weighted sum of Gabor atoms, MAGE permits a more accurate estimate of the amplitude and phase of transient bursts than existing methods. Since Denis Gabor’s pioneering paper on the discrete Gabor Expansion (Gabor, 1946), time-frequency signal analysis has proven to be an important tool for many fields. In neurophysiology, time-frequency analysis has often been used to characterize and describe transient bursts in local field potential data. However, these transient bursts have a wide range of variable durations, suggesting that a time-frequency-scale dictionary composed of elementary signal “atoms” may prove useful to more accurately match recorded bursts. While overcomplete multiscale dictionaries are useful, generating a sparse code over such dictionaries is a difficult computational problem. Existing adaptive algorithms for discovering a sparse description are slow and computationally intensive. Here we describe the Multiscale Adaptive Gabor Expansion (MAGE), which uses an implicit dictionary of parametric time-frequency-scale Gabor atoms to perform fast parameter reassignment to accelerate discovery of a sparse decomposition. Using analytic expressions together with numerical computation, MAGE is a greedy pursuit algorithm similar to Matching Pursuit, restricted to a dictionary of multiscale Gaussian-envelope Gabor atoms. MAGE parameter reassignment is robust in the presence of moderate noise. By expressing a unknown signal as a weighted sum of Gabor atoms, MAGE permits a more accurate estimate of the amplitude and phase of transient bursts than existing methods. A

## I. INTRODUCTION

In neurophysiology, bursts are brief periods of increased neural activity that reflect local network coordination, resulting in transient, non-stationary oscillations that exhibit high trial-to-trial variability. Mounting evidence suggests that transient bursts play an important functional role in neural coding (Lundqvist et al, 2018) and network coordination (Kirst et al, 2016). For example, Lundqvist and colleagues (Lundqvist et al, 2016) showed that the amount of information prefrontal neurons encoded about a stimulus in a working memory task was dependent on narrow-band oscillatory bursts in the gamma band (55-90 Hz). In particular, neurons encoded more information during periods of increased gamma power (greater than 2 standard deviations above the mean spectral power) compared to periods with baseline power levels. In their model, gamma bursts are signatures of cell assembly activation that are associated with informative spiking within single cells. Spikes from single neurons are more informative during bursts (high amplitude or spectral power) compared to inter-burst periods (mean to low power). Furthermore, the spike timing of individual cells relative to the phase of an ongoing high-frequency burst carries information about working memory load (Siegel et al, 2009). Similarly, Battaglia and colleagues showed that the amplitude and phase of burst activity that occurs in multiple areas is associated with the gating and flow of information within a distributed network (Palmigiano et al, 2017). In their simulations, small relative phase differences between bursts had a large impact on the direction and gain of information transmission.

These studies highlight the importance of an accurate estimation of burst parameters, including ongoing amplitude and phase, in order to better understand the role that dynamic bursts play in perception, cognition, and action. In addition, moving from arbitrary experimenter-defined oscillatory bands of interest to a robust, data-driven approach for the identification of transient bursts will help identify what functional role bursts play in computation, communication, and network coordination. However, the detection and accurate estimation of burst parameters is a difficult computational problem.

While a given burst may exhibit a stable instantaneous frequency for the duration of its occurrence, subsequent bursts may exhibit high variability in mean frequency, onset time relative to stimuli, or burst duration. This variability and non-stationarity can make it difficult to study burst dynamics with the traditional methods often used in electrophysiology studies. Traditional methods assume signal stationarity and model the variability of repeated experimental trials as a constant signal waveform embedded in stochastic noise. However, recent studies suggest that the ongoing local field potential (LFP) signal can be thought of as a non-stationary sequence of discrete, variable, and disjoint atomic bursts occurring within a stationary background of relatively low-power continuous ongoing activity. The goal of this paper is to describe a novel adaptive method for detecting transient bursts in non-stationary neurophysiological signals. Specifically, this method describes bursts as Gaussian-envelope Gabor atoms, using parameter reassignment and matching pursuit to generate a sparse atomic decomposition of neural signals. Given an initial set of Gabor parameters describing a burst in terms of time, frequency, scale, amplitude, and phase, this adaptive method returns improved parameters using fast refinement rules based on one-shot curve fitting, similar to the adaptive chirplet method of Yin and colleagues (Yin et al, 2002). Whereas their method used finite differences and 6 inner product operations to estimate the reassigned parameters, this method uses 4 inner product operations computing derivatives with respect to the parameters for time, frequency, and scale (duration in time or bandwidth in frequency). For this reason, we term the method the Multiscale Adaptive Gabor Expansion, or MAGE. In the following we review some of the concepts involved in moving from linear, non-adaptive, and time-invariant signal processing of stationary signals to non-linear, sparse, and adaptive signal processing working on non-stationary signals using overcomplete frames or dictionaries. Next we derive the parameter reassignment rules at the core of MAGE before applying them to simulated and empirically-recorded data examples. The sensitivity and speciificy of MAGE are estimated under noiseless and noisy conditions. Finally, we consider some potential concerns or possibilities for applying MAGE to neurophysiological signals.

### A. From linear, time-invariant processing of stationary signals to sparse, adaptive decomposition of non-stationary signals via overcomplete dictionaries

Signal processing often requires us to characterize an unknown signal *f*(*t*). One way to do so is to compare the unknown signal *f*(*t*) to a dictionary of known references signals *G*, where each element *g_k_*(*t*) ∈ *G* is a time-frequency signal “atom” that captures some feature of interest, such as localization in time, frequency, or scale. The inner product is a useful procedure for directly comparing an unknown target signal *f*(*t*) and a known probe signal *g*(*t*):

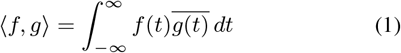

where 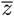 is the complex conjugate of 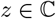.

The more similar *f* is to *g*, the larger the magnitude of the inner product. Conversely, if *f* and *g* are dissimilar, their inner product will tend toward zero. Therefore, the set of inner products of the signal *f* against the dictionary *G* – that is, the set {〈*f, g_k_*〉|*g_k_* ∈ *G*} – describes how similar the unknown signal is to every atom in the dictionary and provides a simple measurement technique for generating structured data about the unknown signal: which reference or probe atoms best describe the unknown target signal, what part of the target signal different atoms capture, and the relative importance of different atoms for accurate and robust signal reconstruction.

A surprising number of signal processing transforms can be characterized in this way, as an ordered set of inner products with a parametric dictionary (Qian and Chen, 1993; Qian and Chen, 1994; Yin et al, 2002). These include 1-dimensional transforms such as the Fourier, Laplace, and fractional Fourier Transforms, 2-dimensional transforms such as the Short-Time Fourier and Wavelet Transforms, and 3-dimensional transforms such as the multi-scale Gabor transform (Cohen, 1995; Grochenig, 2000; Qian, 2001).

#### 1) From orthogonal bases to sparse overcomplete dictionaries

If the dictionary *G* spans the same space as the unknown target signal *f* while each dictionary atom is orthogonal to every other atom (that is, *m* ≠ *n* implies 〈*g_m_, g_n_*〉 = 0 for all *g_m_, g_n_* ∈ *G*), then *G* forms a basis for *f*. In this case, the analysis and synthesis of *f* assumes the form:

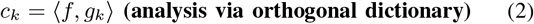

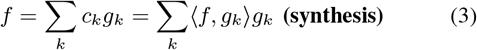

The expansion of a signal over orthogonal dictionaries has proven to be very useful in many domains, most notably in signal compression and communication. However, in other domains such as biomedical imaging, the primary goal is to *characterize* a novel signal, rather than to *transmit* it elsewhere; in this case orthogonal dictionaries may miss signal features that we would like to describe. That is, while an orthogonal basis still provides for perfect reconstruction, describing a single feature of the target signal may require a weighted sum of several different dictionary atoms.

Therefore, for cases where signal characterization is the primary goal, it may be more appropriate to use an *overcomplete* dictionary, also known as a redundant frame. As with a basis, a frame spans the space of the signal – but unlike a basis, frame (or dictionary) atoms need not be orthogonal. Relaxing the orthogonality constraint allows us to include a larger variety of signal atoms in the dictionary, increasing the likelihood that one of the atoms will prove to be a good match to a part of the target signal. As before with the orthogonal dictionary, analysis is simple:

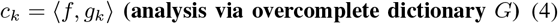

In many cases, such as biomedical research, the goal of signal processing is to generate the set of coefficients {*c_k_*} which are then examined directly for some feature of interest – that is, in many cases there is no need to explicitly reconstruct the original signal *f* from the set of dictionary inner products. If desired, synthesis can still be performed using a synthesis dictionary *G*^†^ (distinct from the analysis dictionary *G*), where *GG*^†^ = *G*^†^*G* = *I*:

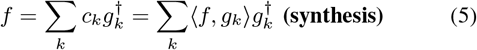

In general, however, synthesis becomes more complicated in the overcomplete setting, in that for a given analysis dictionary *G*, the synthesis dictionary *G*^†^ is not unique.

Alternatively, given a large overcomplete dictionary, one can use an iterative pursuit algorithm to generate a sparse signal description.

#### 2) Sparse Signal Representation via the Matching Pursuit algorithm

Recall that given an overcomplete dictionary *G*, any signal can be described as a sum of weighted atoms:

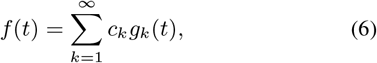

where *m* < *n* implies |*c_m_*| ≥ |*c_n_*|. For greedy algorithms such as Matching Pursuit (Mallet and Zhang, 1993), each coefficient *c_k_* is extracted sequentially. That is, we can express any signal *f*(*t*) as a filtered signal *f_N_* and a residual signal *r_N_*:

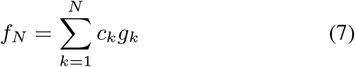

and

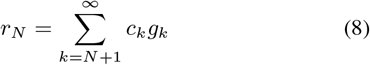

where

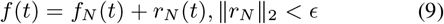

More formally, we can describe the Matching Pursuit algorithm as a function that takes in the unknown signal *s* to be decomposed, the dictionary *D* of known signal atoms, and the number *N* of atoms to be extracted:

#### Matching Pursuit Algorithm [*G*, *c*, *r*] = **MP**[*s*,*N*,*D*] Inputs

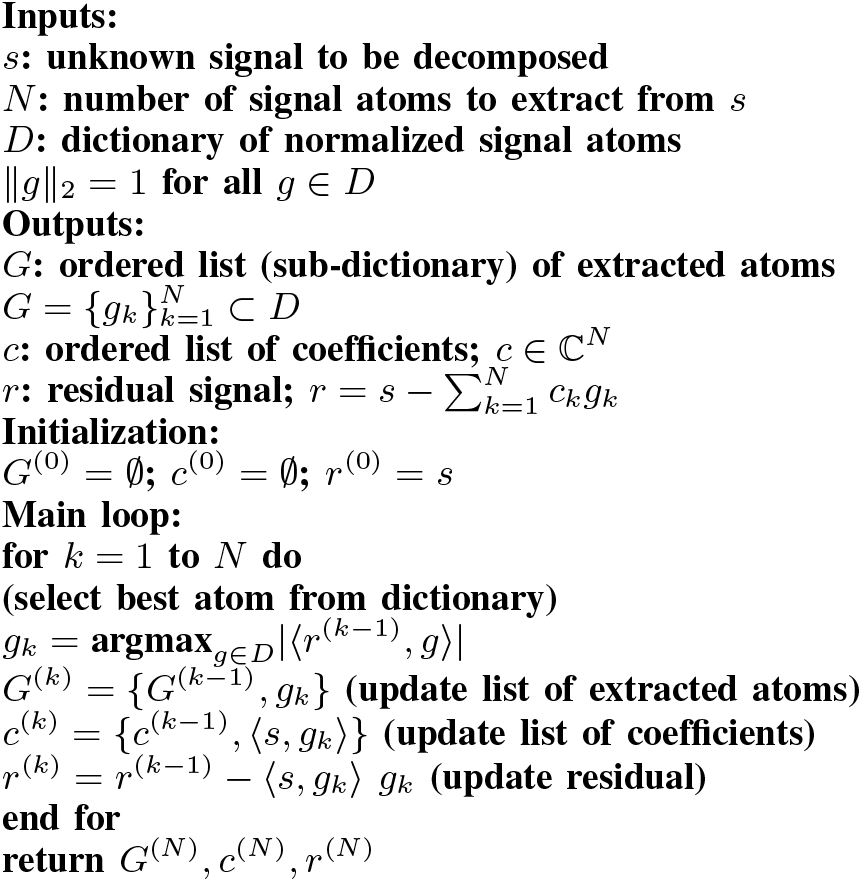

That is, Matching Pursuit finds the next best atom to extract by computing the inner product of the unknown signal *s* with all atoms in the dictionary *D*:

(compute inner product of signal with dictionary atoms)

1. *ρ_γ_* = 〈*r*^(*k*−1)^, *g_γ_*〉 for all *g_γ_* ∈ *D*
2. *γ*\* = argmax_*γ*_|*ρ_γ_*| (find index of best atom)
3. *g_k_* = *g*_*γ*\*_ (update current atom with best atom) (select best atom from dictionary)
4. *g_k_* = argmax_*g*∈*D*_|(*r*^(*k*−1)^, *g*〉|

Unfortunately, steps 1 and 2 are often computatonally expensive. One solution to this computational burden is to restrict the dictionary *D* to only use atoms of a known parametric form, and use the analytic expression for the atom waveform and rather than storing an explicit numerical waveform. Similarly, for some parametric dictionaries the inner product also has an analytic form and can be used to speed up the discovery of the best-fit atom in matching pursuit.

#### 3) The dictionary of multiscale Gaussian-envelope Gabor atoms

One signal atom that is very useful in time-frequency analysis is the Gaussian function *φ*(*t*):

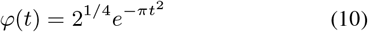

Recall that a signal 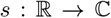 is finite-energy if and only if

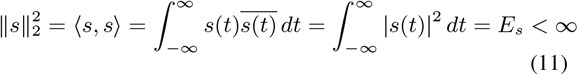

Direct integration shows that *φ* has unit energy,

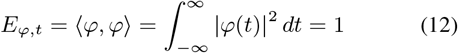

We can use the Gaussian energy density function to compute the average time of the signal:

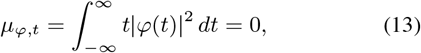

Similarly, we can compute the time-domain variance:

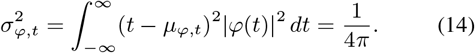

Recall the definition of the Fourier transform 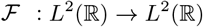 and its inverse 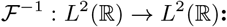:

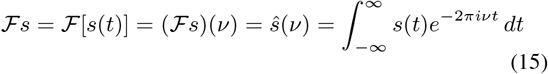

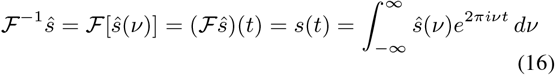

which we take here as already extended to be bounded, linear, unitary operators. Applying the Fourier Transform to the time-domain Gaussian function, we find the frequency-domain expression

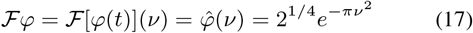

From this we determine the frequency-domain energy, mean frequency, and frequency-domain variance:

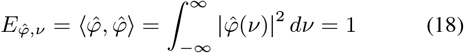

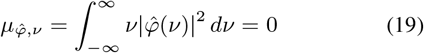

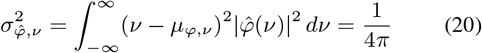

Here we see why Gaussian functions are so useful in time-frequency analysis. Recall that for any signal *s*,

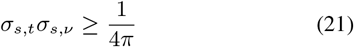

meaning that there is a lower limit on how concentrated a signal can simultaneously be in both the time- and frequency-domains. In general, if the energy of *s* is concentrated in time, then the energy of *s* is spread out in frequency, and vice versa. However, Gaussian-type functions make this inequality an equality; that is,

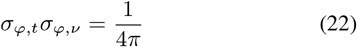

meaning that Gaussian functions are maximally localized in time and frequency. There exist no signals which have a tighter concentration of energy in the time-frequency plane. Therefore, if you want to decompose an unknown signal by describing in terms of simple, known time-frequency “atoms,” then Gaussian functions are an excellent choice, given their strong energy concentration.

#### 4) The translation, modulation, and dialation operators

To be able to move *φ* to any location in the time-frequency plane, we introduce two operators 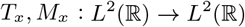, where 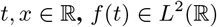.

The translation operator

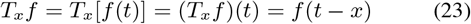

shifts or translates the signal *f* in time, while the modulation operator

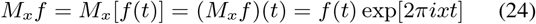

shifts or modulates the signal *f* in frequency. Applying these operators to *φ* permit us to shift the mean time and mean frequency of the localized signal atom. That is, given the simple reference function *φ* as a fundamental signal “atom” which is well-localized in time and frequency, together with the operators *T_x_* and *M_x_* that move signal atoms in time and frequency, we can fully tile an unknown signal within the time-frequency plane, comparing small segments of this unknown signal to time- and frequency-shifted version of the known reference signal *φ*. To describe an unknown signal *f*, we can compute the inner product of it with the dictionary 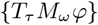. This is the origin of the continuous STFT (where 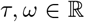) as well as its sampled counterpart, the Gabor expansion (where *τ* = *am* and *ω* = *bn* for small, fixed *a, b* ∈ ℝ and 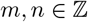).

However, time- and frequency-shifts alone do not capture all signal features of interest. Complex signals such as the local field potential recorded in neuroelectrophysiology often involve components that reflect different durations or scales – some signal sub-components may rise and decay quickly while others evolve more slowly.

We can examine variations in scale by introducing another operator *D_x_* : *L*^2^(ℝ) → *L*^2^(ℝ), where the dialation operator

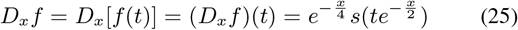

dialates or re-scales the input domain of *f* while preserving the signal energy.

Given the Gaussian function *φ* together with the translation, modulation, and dialation operators defined above, we can generate an overcomplete dictionary *G* ={*T_a_M_b_D_c_φ*|*a,b,c* ∈ ℝ^3^}. This dictionary tiles the time-frequency plane, in that for each time-frequency coordinate (*a, b*) we can find an atom centered at that location – in fact, centered at each time-frequency point (*a, b*) is a continuous family of atoms that span all possible scales or durations specified by the duration parameter *c*. Furthermore, since the Gaussian *φ*satisfies the uncertainty principle relation as an equality, each such atom provides the most compact time-frequency energy concentration possible.

Specifically, let

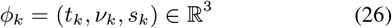

represent the (time, frequency, scale) parameters for a given atom *g_k_* ∈ *L*^2^(ℝ). Then

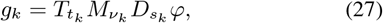

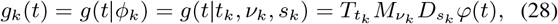

such that

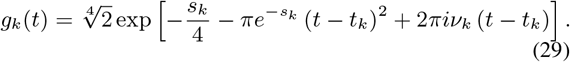

Applying the Fourier Transform to *g_k_* results in

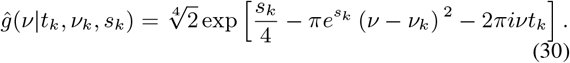

Given these time- and frequency-domain formulas of the atom *g_k_*, we can calculate the total energy, the mean time, and the mean frequency of the Gabor atom:

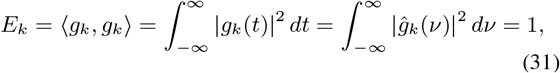

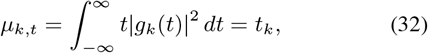

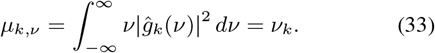

This shows that the generative parameters (*t_k_*, *ν_k_*) correspond to the measured values of mean time and mean frequency.

Finally, as a measure of signal duration, we can calculate the time-domain and frequency-domain variance:

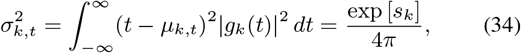

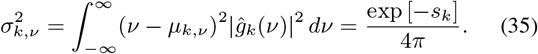

Showing that the duration parameter *s_k_* captures the variance in signal energy in both the time and frequency domains.

#### 5) Inner product relation between Gabor atoms

Using the parametric formula for *g_k_* given above, we can identify an analytic expression for the inner product between two Gabor atoms:

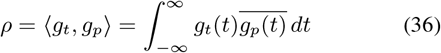

such that

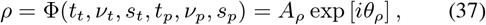

where

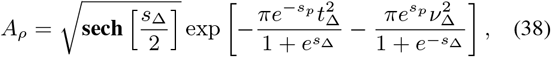

and

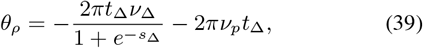

where the difference between target and probe Gabor parameters are given by

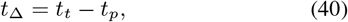

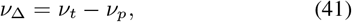

and

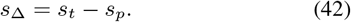

Thus, the inner product between two gabors can be computed numerically or analytically. This is useful once we recall that, as in Matching Pursuit, any signal can be decomposed into a weighted sum of atoms and a residual signal:

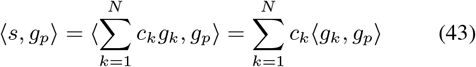

Since the derivate operator is conjugate linear, we also have

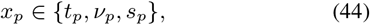

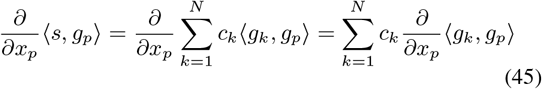

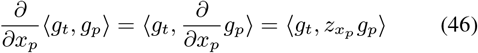

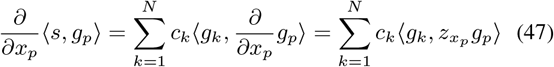

for some *z_x_p__*. To compute *z_x_p__*, recall that the derivative of the inner product of two gabors with respect to a given Gabor parameter is equal to the inner product of a gabor atom with a derivative atom. That is, we can exchange the derivative and inner product operators.

#### 6) Derivation of MAGE parameter reassignment rules

Recall that our goal is to derive parameter reassignment rules such that if we are given a coarse parameter estimate of a target Gabor atom, then our reassignment rules will give us the parameters for a Gabor atom nearer to the target (ideally in one step). Since the operators in the prior sections are conjugate linear, we can exchange the order of operators to generate analytic expressions that are easier to deal with, as when we replace the derivative of an inner product between two Gabor atoms with the inner product between a target Gabor and a derivative kernel. Finally, we note that these derivative atoms have an analytic form, allowing us to rearrange and simplify to produce our reassignment rules.

Note that the 1st derivative of probe Gabor atom with respect to the center time parameter *t_p_* is

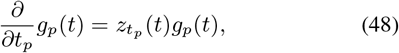

where

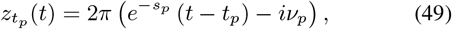

The 1st derivative of probe Gabor atom with respect to center frequency parameter *ν_p_* is

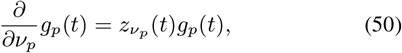

where

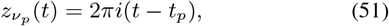

The 1st derivative of probe Gabor atom with respect to the duration parameter *s_p_* is

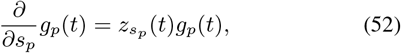

where

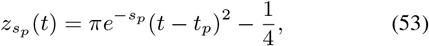

Let *m* stand for numerically-measured inner product values. Note that

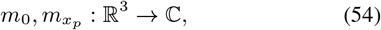

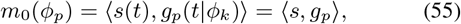

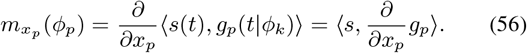

Analytically, we have

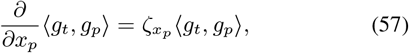

for

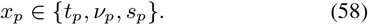

and

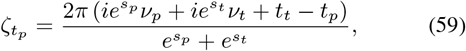

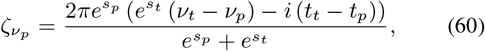

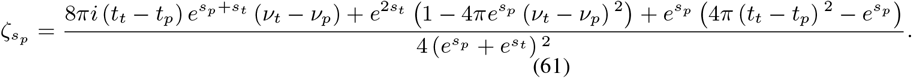

These terms allows us to define a reassignment function for each Gabor parameter:

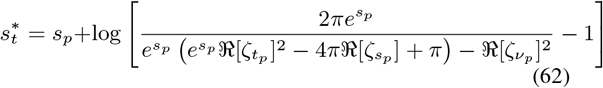

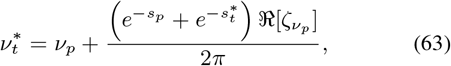

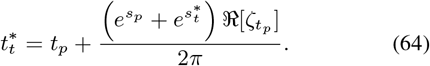

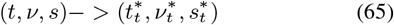

where 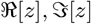 stand for the real and imaginary parts of a complex number z.

Interestingly, not all derivatives are needed, since the derivative of the inner product of two Gabor atoms with respect to center time and center frequency are related, with the imaginary part of one correponding to the real part of the other. That is, the relation of *ζ_t_p__* to *ζ_ν_p__* is:

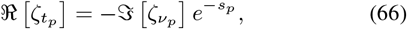

and

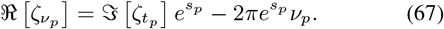

Thus, given an initial probe Gabor *g_k_* that has a moderate inner product magnitude with an unknown signal *s*, MAGE provides parameter reassignment formulas that may represent a Gabor with a better fit to the unknown signal. Given a moderately good initial probe Gabor, the reassignment function 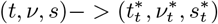 suggests the next test point in time-frequency-scale space.

#### 7) From Matching Pursuit to the Multiscale Adaptive Gabor Expansion (MAGE)

This gives us an update function for parameter reassignment which we can include within the Matching Pursuit algorithm to produce MAGE:

#### Multiscale Adaptive Gabor Expansion Algorithm

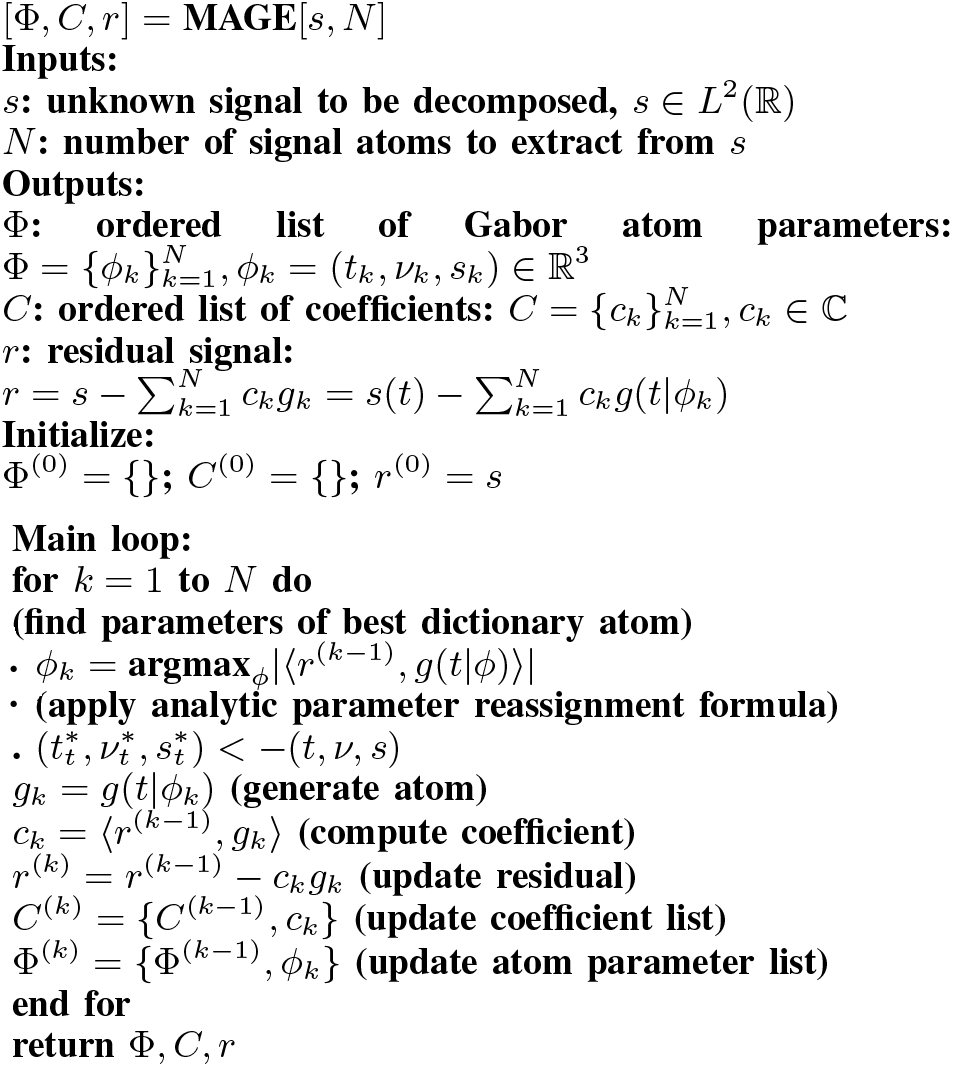

Including parameter reassigment rules within Matching Pursuit, as we do to develop MAGE, is justified for several reasons. First, recall that most matching pursuit algorithms use an explicit dictionary of sampled waveforms, such that Matching Pursuit only explores a discrete subset of parameter values corresponding to dictionary atoms. MAGE, however, performs its parameter reassignment in a continous parameter space relatively unaffected by issues of sampling or regularity of the search grid (eg, rectangular vs. triangluar-quincunx). This means that initial sampling grid can be fairly coarse to speed up the numerical computation, while relying on the MAGE parameter reassignment to generate a more refined estimate. Second, the analytic Gabor inner product relation used in the derivation of the reassignment rules can be used to quickly determine if a Gabor inner product value needs to be updated or not, greatly speeding up the Matching Pursuit algorithm. Third and finally, multiple probe Gabors (with different specific noise interference terms) can be used to estimate the parameters for the same target Gabor, implementing a simple form of noise reduction.

### B. Example application of MAGE to synthetic target signal with additive noise

Here we provide some examples of use to show how the MAGE parameter reassignment rules may assist in time-frequency-scale signal analysis. First, we highlight some issues with traditional linear FIR filtering, showing that adaptive, non-linear techniques such as Matching Pursuit overcome many of these difficulties. We then show that the MAGE reassignment rules increase the effective resolution of a Matching Pursuit search grid at a small computational cost.

While linear finite-impulse-response (FIR) filtering is fast and has proven very useful over the past several decades, there are several reasons why researchers engaged in non-real-time analysis should consider using adaptive nonlinear techniques such as Matching Pursuit and MAGE.

First, traditional approaches in neurophysiology have often employed pre-selected bands (e.g., analyzing theta-, beta-, and gamma-band activity). Non-overlapping bands and coarse/sparse filterbanks risk missing the occurence of bursts that occur in-between the FIR filters used, and will de-emphasize bursts that occur on band edges. In contrast, adaptive and data-driven techniques such as Matching Pursuit and MAGE use a formal definition of signal energy rather than predefined oscillatory bands to identify segments of transient burst activity, and are therefore less likely to result in signal detection MISSES due to lack of filter coverage.

Second, dense overlapping filterbanks may register one actual burst in two or more filter channels, resulting in a signal-detection FALSE ALARM. In contrast, adaptive techniques such as Matching Pursuit and MAGE add a non-linear winner-take-all inhibition process to prevent one atom from appearing more than once in an atomic decomposition. That is, a single burst appearing in two overlapping bands requires additional processing to remove false signals and preserve signal energy.

Third, there are many reasons to expect signals to be composed of independently generated bursts that overlap in time, frequency, and scale. Signals that differ slightly in frequency and scale will produce interference terms, and may lead to the improper estimate that one burst is present when there are actually two or most bursts in the same time-frequency-scale region in parameter space. Sparse representations of dense signals is an active area of signal processing research, but it is known that linear FIR filtering alone will misidentify this situation. For example, suppose neural populations A and B are spatially intermixed, but in terms of connectivity are essentially separate networks. A generates short-duration bursts around 35 Hz while B generates long-duration bursts around 45 Hz. Field potential recordings will observe a weighted linear combination of these two distinct bursts. While an array of spatially-separated electrodes may be used to solve this source-separation problem, sparsification of mixed signals is not a straightforward problem.

Fourth, lossless energy-preserving transforms that permit analysis and synthesis to be performed at any stage of processing is to be preferred to analysis techniques that are lossy and do not permit recovery of the original signal. Only attending to pre-selected bands will result in the loss of information either through missing information (problem 1 above) or using non-orthogonal filters (problem 2 above). In general, analysis and synthesis kernels are not the same (Qian, 2001). Adaptive techniques such as Matching Pursuit and MAGE can employ the same kernel (here, Gaussian-weighted Gabor atoms) by employing a greedy non-linear selection process.

For the four reasons given above, researchers currently using linear FIR filtering may want to consider using adaptive, data-driven techniques such as Matching Pursuit and MAGE. These techniques are slower than linear FIR filtering alone, but have several advantages over non-adaptive techniques. One way to conceptualize the MAGE parameter reassignment rules is that it effectively provides a denser search grid for decomposition at similar runtimes and computational cost. That is, to get the same SNR decomposition, one can use Matching Pursuit with a dense search grid (cf Fig 1B) or use a coarse search grid together with MAGE parameter reassignment. By exploiting knowledge about the analytic form of Gabor atoms, the derivative atoms with respect to the defining parameters, and the inner product relation between distinct Gabor atoms, the reassignment rules effectively provide Matching Pursuit with a dense, implicit dictionary. Figure 5A shows that the run-time for Matching Pursuit and MAGE increases with the density of the initial parameter space search grid, but that there is little difference between the two algorithms in runtime for the same grid density. In contrast, Figure 5B shows the SNR for Matching Pursuit and MAGE as a function of grid size, while Figure 5C shows the SNR as a function of runtime. For a given initial grid size, including MAGE reassignment will improve your results with a negligible runtime cost. Alternatively, for a given SNR, MAGE will take less time to produce this decomposition than Matching Pursuit alone. Given the apparent utility of the analytic MAGE parameter reassignment rules, we consider two examples in more detail below.

**Fig. 1:**
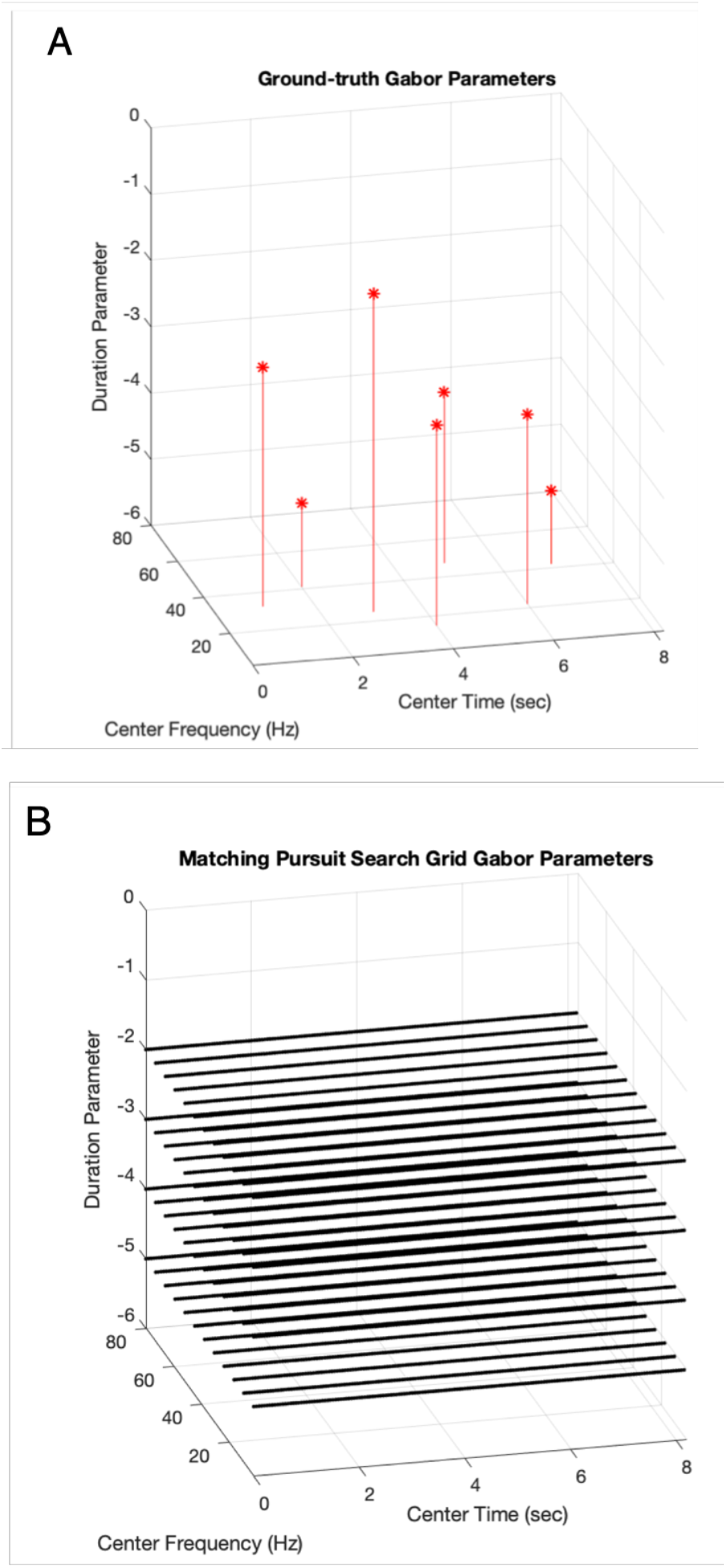
Three-dimensional time-frequency-scale parameter space for Gabor atoms. A) Example parameters (red) for the seven Gabor atoms of a simulated target signal. Each Gabor atom is defined by three parameters of center time ttact, center frequency vtact, and duration parameter stact together with scalar amplitude and phase values. B) A 12×8192×4 search grid of Matching Pursuit Gabor atoms, with four scale levels, twelve center frequencies, and 8192 center times. Note that the parameters for most target atoms will not fall on the Matching Pursuit search array. The purpose of MAGE is to refine the parameters for the best Matching Pursuit Gabor atom estimate, by reassigning the best-estimate MP parameter tuple (tp,vp,sp) for a probe Gabor atom to the parameter tuple (ttest, vtest, stest) for the estimated target Gabor atom.

#### 1) Example of MAGE use: synthetic data with known ground-truth under additive noise

Figure 1 shows an example where MAGE is applied to a synthetic signal where there is a known ground-truth. Figure 1A shows the parameters for seven weighted Gabor atoms, and Figure 1B shows the initial Matching Pursuit search grid of 12 center frequencies, 4 scale levels, and 8192 center times. This is efficiently computed as convolution of the signal with 48 (12×4) filters in a filterbank. The element of the sampling grid with the strongest inner product with the largest target atom is selected after this initial population of the search grid. Note that in general the parameters of target atoms do not fall exactly on the Matching Pursuit search grid.

In contrast to Figure 1 which deals with a 3-D (time, frequency, scale) parameter space, Figure 2A (signal space) shows real component of the synthetic signal thus generated (red) along with the synthetic signal plus additive noise (black). The signals are complex-valued functions of time over some support from initial time to final time. The center time of each Gabor atom is within this signal time support. Figure 2B shows preselected FIR filtered versions of noisy synthetic signal in the theta, beta, and gamma bands. Note that some atoms appear in multiple bands, highlighting the need for an inhibitory winner-take-all process to prevent FALSE ALARMS. A single atom is highlighted in Figures 2C-E, with the ground truth target signal shown in red, and the burst estimates provided by linear FIR filtering (Figure 2C), by Gabor-based Matching Pursuit without MAGE parameter reassignment (Figure 2D), and by MAGE (Figure 2E). Note the mismatch between target burst and the best coarse-grid Matching Pursuit estimate; this mismatch is what the non-iterative, analytic MAGE reassignment rules are designed to overcome.

**Fig. 2:**
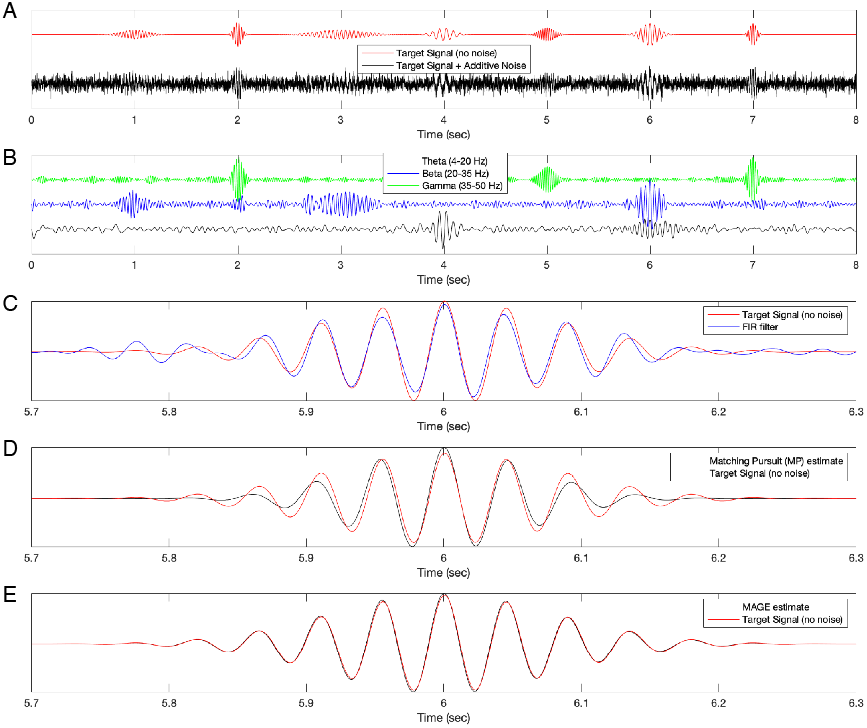
Example of transient oscillatory bursts and burst identification. A) Target signal with 6 gabor atoms with varying center frequencies and durations (red), and target signal with additive noise. B) Target signal filtered withl inear FIR filter in the theta (4-20 Hz), beta (20-35 Hz), and gamma (35-50 Hz) frequency bands. C) Example of ground-truth burst (red), and the estimated burst (blue) extracted by linear FIR filtering plus thresholding. D) As in C, but for atom extracted by adaptive, non-linear matching pursuit. E) As in C, but for atom extracted by MAGE parameter reassignment, which refines a coarse estimate provided by matching pursuit.

Figure 3 shows a comparison of the estimated Gabor atoms output by Matching Pursuit and MAGE for this known-ground-truth synthetic signal. Both Matching Pursuit and MAGE identify all atoms, but the inner product between actual target Gabors and estimated target Gabors tends to be larger for MAGE than for Matching Pursuit without parameter reassignment. Both algorithms extract the atoms sequentially based on burst energy, with each extraction reducing the energy of the residual signal and improving the SNR of the filtered signal estimate (Figure 3C).

**Fig. 3:**
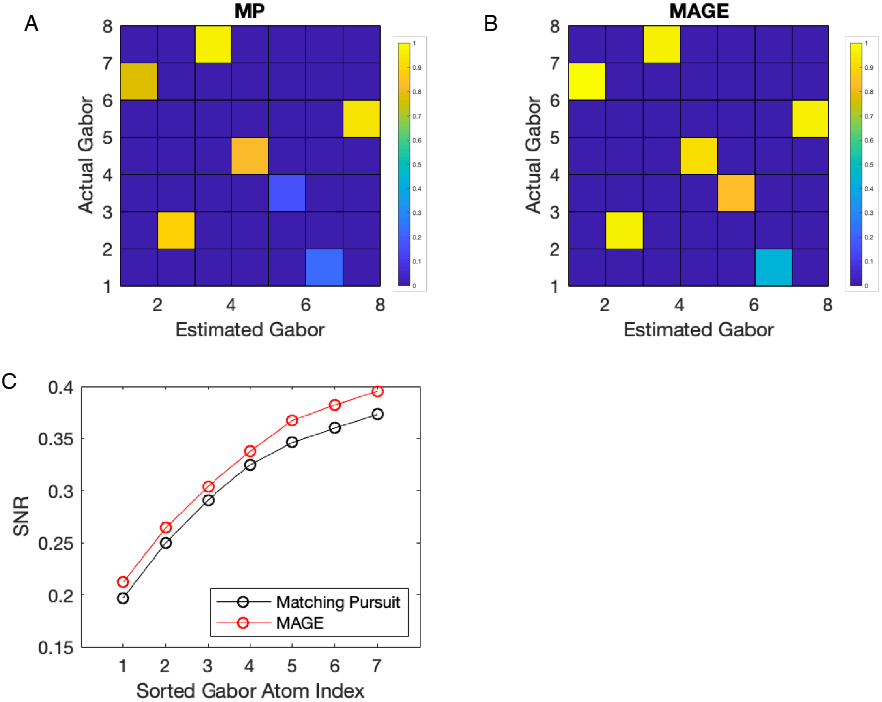
Confusion matrix for actual-estimated Gabor atoms generated by Matching Pursuit and MAGE. A) Inner product magnitude between the estimated target Gabor atoms generate by Matching Pursuit without reassignment (MP) and actual Gabor atoms. B) As in A, for MAGE parameter-reassigned estimated target Gabor atoms. C) Signal-to-noise increase due to repeated extraction of strongest Gabor atoms, as found by Matching Pursuit (black) and MAGE (red).

### C. Sensitivity and specificity under noiseless and noisy conditions

Figure 4 provides information about the influence of noise on the MAGE parameter reassignment rules. Given noise, a single target Gabor atom (actual target), and a initial probe Gabor (estimated target) with a large degree of error (inner product between actual target Gabor and probe estimate is 0.2, indicating small overlap), MAGE reassignment performs well under noiseless or low noise conditions (Figure 4A). Note that without noise, MAGE parameter reassignment moves the probe Gabor from an inner product magnitude of 0.2 to ¿0.95 in one step, indicating that the target estimate is a good match to the actual target. Here we define an actual-estimate target Gabor inner product magnitude as a HIT when it exceeds a threshold of 0.95 (colorless area in Figure 4A). For 256 distinct target atoms at each noise level, 128 probe estimates with an actual-target/initial-probe inner product magnitude of 0.2 (initial) were used to estimate a final probe atom (target estimate). Figure 4A shows the average inner product magnitude between the actual-target/estimated-target Gabors after application of the MAGE parameter reassignment rules. For clarity, for each noise level the inner product values were sorted. As noise increases, the average final inner product value drops below the HIT inner product threshold, but in most cases MAGE reassignment results in an increase of inner product magnitude. Figure 4B collapses across these individual probes to show the average improvement in the final target estimate (blue) compared to the initial target estimate (red dotted). For a fraction of cases (approximately 15%) under strong noise conditions, the final probe estimate is worse than the inital estimate. This reassignment failure is likely due to constructive and destructive interference between Gabor atoms changing the energy landscape and violating the sparsity assumptions (no overlapping bursts) used in the derivation of the reassignment rules. Formulating new reassignment rules for multiple overlapping bursts may be possible due to the linearity of the operators involved, but will likely involve several more sample points than the sparse-signal MAGE reassignment rule considered here.

**Fig. 4:**
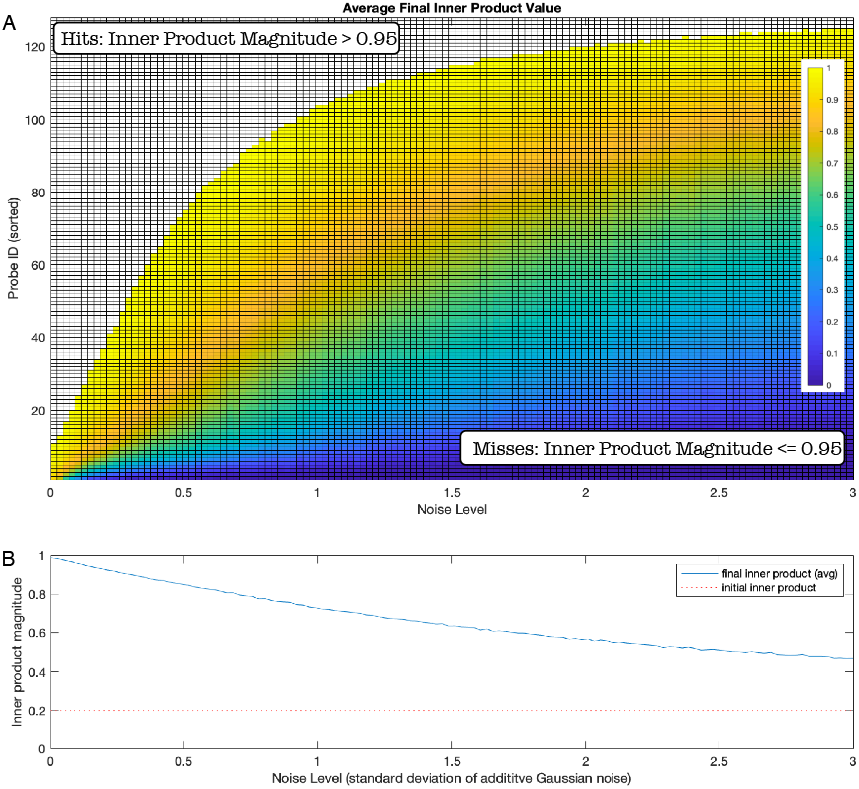
Noise analysis for MAGE parameter reassignment. A) For a given noise level and target Gabor atom gt, A probe Gabor gp with a target-probe Gabor atom inner product magnitude value of 0.2 is reassigned by MAGE. Hits are considered reassigned Gabor atoms that have an inner product magnitude of 0.95 or greater. As noise increases, the inner product magnitude between estimated and actual target Gabor atoms drops, with a higher fraction of misses, but improved upon the initial MP estimate. B) Probe Gabor atoms that have an initial probe-target inner product value of 0.2 (red line) have their parameters reassigned to an estimated target Gabor atom. At low noise levels, reassigned parameter estimates generate final inner product values near 1 (noiseless). As noise increases, the accuracy of the probe Gabor parameter reassignment drops (blue line).

**Fig. 5:**
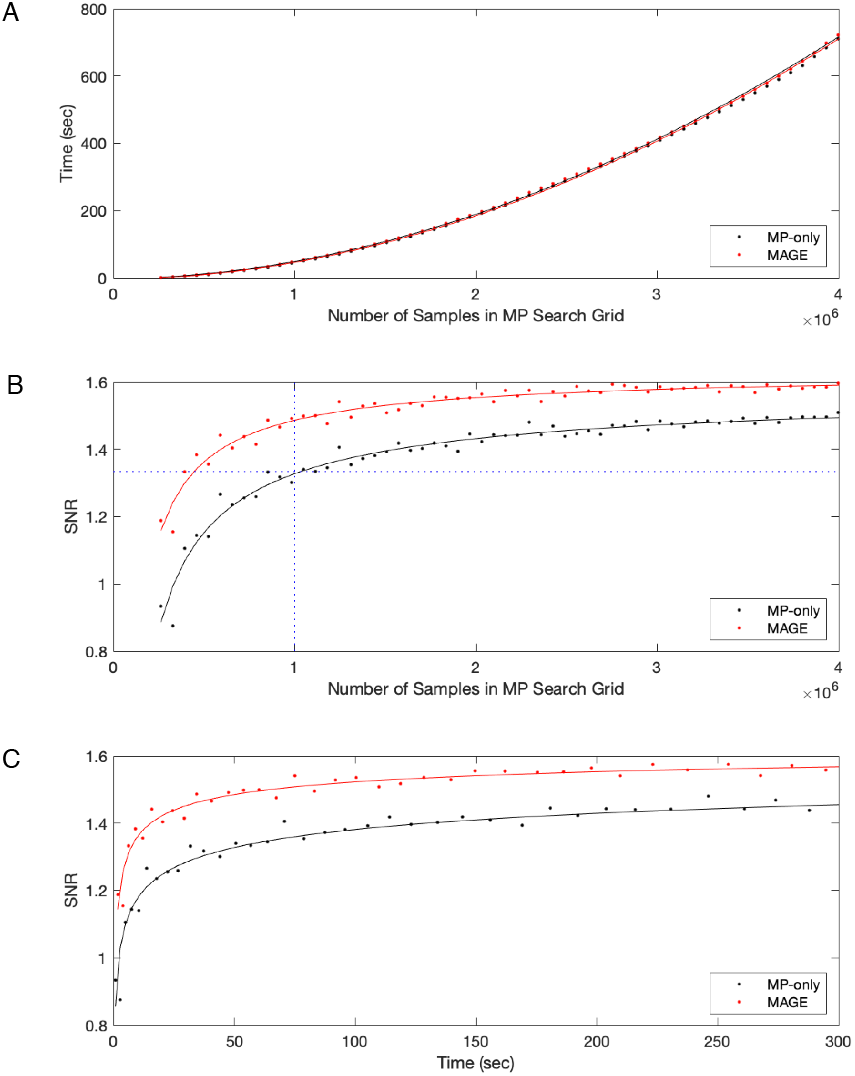
Performance of MAGE and Matching-Pursuit-only (no one-step parameter reassignment). A) For both Matching Pursuit (MP) and MAGE, a coarse sampling grid (left) leads to a faster runtime than a dense sampling grid (right). However, the inclusion of MAGE parameter reassignment does not lead to a significant increase in runtime as compared to Matching-Pursuit-only run on an identical sampling grid. B) As the density of the sampling grid increases, the SNR of the extracted signal increases for both MAGE and MP. As shown by the vertical dotted line (blue), the SNR produced by MP at a specific sampling grid density can be achieved with a coarser grid if MAGE reassignment is included. Alternatively, for a given sampling grid density, MAGE consistently produces better results in terms of SNR of the extracted signal. C) The SNR as a function of runtime for both MAGE and MP. Given a specific runtime and SNR on the MP curve, MAGE can generate an equivalent SNR with a reduced runtime. Alternatively, for a specific fixed runtime, MAGE will output results with a higher SNR than MP.

### D. Discussion and Conclusion

While the theoretical foundations of Matching Pursuit and similar adaptive algorithms is well established (Mallet and Zhang, 1993; Qian and Chen, 1994), fast and practical numerical implementations remain an open research topic. One of the most useful options right now is the Matching Pursuit Toolkit or MPTK (Krstulovic and Gribonval, 2006), a C++ open source toolkit which maintains a list of atoms that are local maxima in amplitude space and updates only those atoms that have a large inner product with the last atom extracted. MAGE can be seen as a similar augmentation of Matching Pursuit, where the known analytic expressions for Gaussian-envelope Gabor atoms is used to estimate the location of the best local maximum in parameter space, even if it is off of the Matching Pursuit search grid. That is, Matching Pursuit finds the best Gabor atom in its set of regularly sampled atoms (initial coarse estimate step in sampled discrete space), which is then input into the MAGE reassignment rules to find an improved estimate (fast parameter refinement step in continuous parameter space). This fast refinement is a non-iterative parameter reassignment, although an iterative version of MAGE may be useful for denoising. Since MPTK partitions its search tree using time alone (no need to update atoms that do not overlap), it would be interesting to see if MAGE could be incorporated into MPTK, since the inner product relation between any two Gabor atoms can be used to partition the search space in time, frequency, and scale rather than just time alone.

Another concern is the relation between empirically-observed bursts and the parametric model of Gabor time-frequency-scale atoms. Gabor atoms are essentially Gaussian-windowed sinusoids, but recent studies have reminded researchers of the often non-sinusoidal nature of empirically-recorded electrophysiological brain signals (Cole and Voytek, 2017). One response to this concern is to note that the mean time and mean frequency are well-defined concepts for any finite-energy signal (Cohen, 1994). Recall that the center time and center frequency parameters correspond to the mean time and mean frequency, respectively, of any Gabor atom. Similarly, the time-domain variance and the frequency-domain variance of any finite-energy signal can be estimated from the numerical vector representing a sampled version of it, and these values can be used to estimate the Gabor atom scale parameter. So up to the level of 2nd order signal statistics, parametric Gabor atoms and non-parametric (empirically-recorded) signals can be matched. Additionally, data-driven learning methods such as single-channel Independent Component Analysis (ICA; Bell and Sejnowski, 1995) and Kernel-Singular Value Decomposition (k-SVD; Elad, 2010) can be used to further refine a non-parametric description of a burst once the best-fit parametric Gabor atom has been identified. For example, Brockmeier and Principe (2016) show that alternating Matching Pursuit with data-driven kernel-learning via k-SVD and ICA generate a small set of waveforms with consistent clustering properties that permit experimenters to express human EEG via a sparse code. In this regard, MAGE could be used as an initial step in data-driven kernel learning techniques used to identify segments of raw data containing high-power bursts, thus making learning more efficient.

The goal of this paper was to introduce an improved version of fast parameter refinement, similar to that suggested by [Yin et al, 2002]. As with their algorithm, MAGE is performing a curve-fitting operation in order to suggest an atom with a better fit to a signal to be filtered. The fast refinement algorithm of Yin et al uses finite differences to compute points for curve fitting. The spacing between points is an arbitrary hand-tuned parameter which MAGE does away with by using derivatives. Importantly, these derivatives are used to compute values for curve fitting, and are not used for an iterative hill-climbing optimization. Also, while the Yin algorithm uses 6 points for curve fitting, MAGE uses 4 points, which reduces the number of multily-add operations required. Thus, we see the MAGE algorithm as an incremental improvement to the novel agorithm of Yin et al (2002) which may prove useful to researchers in multiple fields.

All MATLAB and Mathematica code available at https://github.com/rcanolty/MAGE

